# Stability and dynamics of a spectral graph model of brain oscillations

**DOI:** 10.1101/2021.12.02.470983

**Authors:** Parul Verma, Srikantan Nagarajan, Ashish Raj

## Abstract

We explore the stability and dynamic properties of a hierarchical, linearized, and analytic spectral graph model for neural oscillations that integrates the structuring wiring of the brain. Previously we have shown that this model can accurately capture the frequency spectra and the spatial patterns of the alpha and beta frequency bands obtained from magnetoencephalography recordings without regionally varying parameters. Here, we show that this macroscopic model based on long-range excitatory connections exhibits dynamic oscillations with a frequency in the alpha band even without any oscillations implemented at the mesoscopic level. We show that depending on the parameters, the model can exhibit combinations of damped oscillations, limit cycles, or unstable oscillations. We determined bounds on model parameters that ensure stability of the oscillations simulated by the model. Finally, we estimated time-varying model parameters to capture the temporal fluctuations in magnetoencephalography activity. We show that a dynamic spectral graph modeling framework with a parsimonious set of biophysically interpretable model parameters can thereby be employed to capture oscillatory fluctuations observed in electrophysiological data in various brain states and diseases.

## 1 Introduction

One of the prevailing questions in the field of neuroscience is to understand brain functional activity, its relationship with the structural wiring, and its dynamics within a brain state and among different brain states and diseases [1–4]. These questions are being pursued using various non-invasive neuroimaging modalities to measure the functional activity and the anatomical structural wiring of the brain. Functional activity is measured using modalities such as functional magnetic resonance imaging (fMRI), electroencephalography (EEG), and magnetoencephalography (MEG) that capture different spatial and temporal scales. The gray matter structural wiring of the brain is estimated with MRI followed by using diffusion tensor imaging (DTI) and tractography. Subsequently, data-driven and model-based approaches are used to understand how structural-functional (SC-FC) relationships arise in different brain states and diseases. Both of these approaches are largely based on transcribing the brain anatomy into a graph, where different brain regions are the nodes connected to each other as edges made of the white matter fiber.

Data-driven approaches have been broadly based on calculating graph theoretic measures to derive SC-FC relationships [5–18, 1]. However, such approaches suffer from a lack of neural physiology [19] and in their inability to infer mechanistic insights. Most modeling approaches have been primarily based on either extensive non-linear models such as neural mass and mean field models [20–30] that require time consuming simulations, or have been linear (based on network control theory) that exclude the biophysical details of the excitatory and inhibitory neuron population ensembles [31–34]. A modeling approach lying between these two extremities was only recently demonstrated by a linear biophysical model called the spectral graph theory model (SGM) that can accurately capture the wide-band static frequency spectra obtained from MEG. This model incorporates biophysics while maintaining parsimony and requiring minimal computation speed [35].

In this paper, we investigate the properties of SGM and extend it to capture the temporal fluctuations in MEG activity, an emerging marker of brain function [36–44]. We focus on SGM for various reasons. Firstly, SGM yields a closed-form steady state frequency response of the functional activity generated by fast brain oscillations. It is based on eigen-decomposition of a graph Laplacian, drawn from the field of spectral graph theory [45–48]. It is a hierarchical model consisting of excitatory and inhibitory population ensembles at the mesoscopic level, and excitatory long-range connections at the macroscopic level. The key distinguishing factor of SGM is that it captures SC-FC using a parsimonious set of biophysically interpretable model parameters: the neural gains, time constants, conduction speed, and macroscopic coupling constant. Due to its parsimony and its ability to directly capture the wideband frequency spectra, parameter inference is more tractable as compared to the non-linear modeling approaches.

Despite prior success of the SGM’s ability to fit wideband regional power spectra using a closed-form analytical solution [35], its ability to accommodate more complex dynamics including regimes of stability and instability have not yet been explored. Other aspects of SGM’s biological relevance remain unaddressed. For instance, it is not known whether a linear model like SGM can accommodate dynamic changes in model parameters that may then lead to dynamic complex behavior. Since the SGM was formulated in terms of steady-state frequency spectra, its transient behavior was not previously addressed. These are important questions, since a biologically realistic model requires sufficiently rich temporal dynamics at all time scales, and stationary spectral response only tells part of the story.

In this paper we address these aspects. Using a series of analytical and numerical explorations, we show that SGM is capable of generating frequency rich spectra and qualitatively different solution regimes in the time domain. In particular, we demonstrate that this model can exhibit combinations of different dynamical solutions: damped oscillations, limit cycles, and unstable oscillatory solutions, depending on the parameter values. We further demonstrate that the dominant alpha band behavior is independent of local oscillators, and can arise purely from the macroscopic network. We then performed a stability analysis to identify stability boundaries separating these different dynamical regimes. In contrast to prevailing dynamic function studies based on noise-driven fluctuations around a bifurcation point of non-linear SC-FC models, we employed SGM to capture temporal fluctuations in the frequency spectra. We show a novel approach to capture dynamic function with only a small set of time-varying model parameters: the neural gains and the macroscopic coupling constant.

## 2 Results

The model used here is a modified SGM we developed recently [49]. We hierarchically model the local cortical mesoscopic and long-range macroscopic signals for each brain region, where the regions are obtained using the Desikan-Killiany atlas [50]. We then solve the model equations to obtain a closed-form solution in the Fourier frequency domain. This provides us frequency rich spectra which is an estimate of the source-reconstructed MEG spectra. At the mesoscopic level, the model parameters *g*_ee_, *g*_ii_, *g*_ei_, are neural gain terms for the excitatory, inhibitory, and population that alternates between excitatory and inhibitory, respectively, and parameters *τ*_e_, *τ*_i_ are characteristic time constants. At the macroscopic level, the parameters are macroscopic graph time constant *τ*_G_, global coupling constant *α*, and conduction speed *v.*

### 2.1 SGM can exhibit oscillations that are damped, limit cycles, or unstable

We explored the transient behaviour of the model by simulating the model solution in the time domain. First, in order to obtain time simulations, we performed a numerical inverse of the Laplace transformed equations, explained in the Methods section. We performed this first for the local mesoscopic model alone and then for the complete macroscopic model, with and without the input Gaussian white noise. The simulations of the mesoscopic model are shown in Fig. 1. As shown in Fig. 1B, three types of solutions are possible: damped oscillations, limit cycles, and unstable oscillatory solutions, depending on the values of the model parameters. Such damped oscillations and limit cycles are observed in the nonlinear neural mass model as well [51]. Upon inclusion of noise in Fig. 1A, damped oscillations and limit cycles are not distinguishable. However, oscillations are observed regardless.

**Figure 1:**
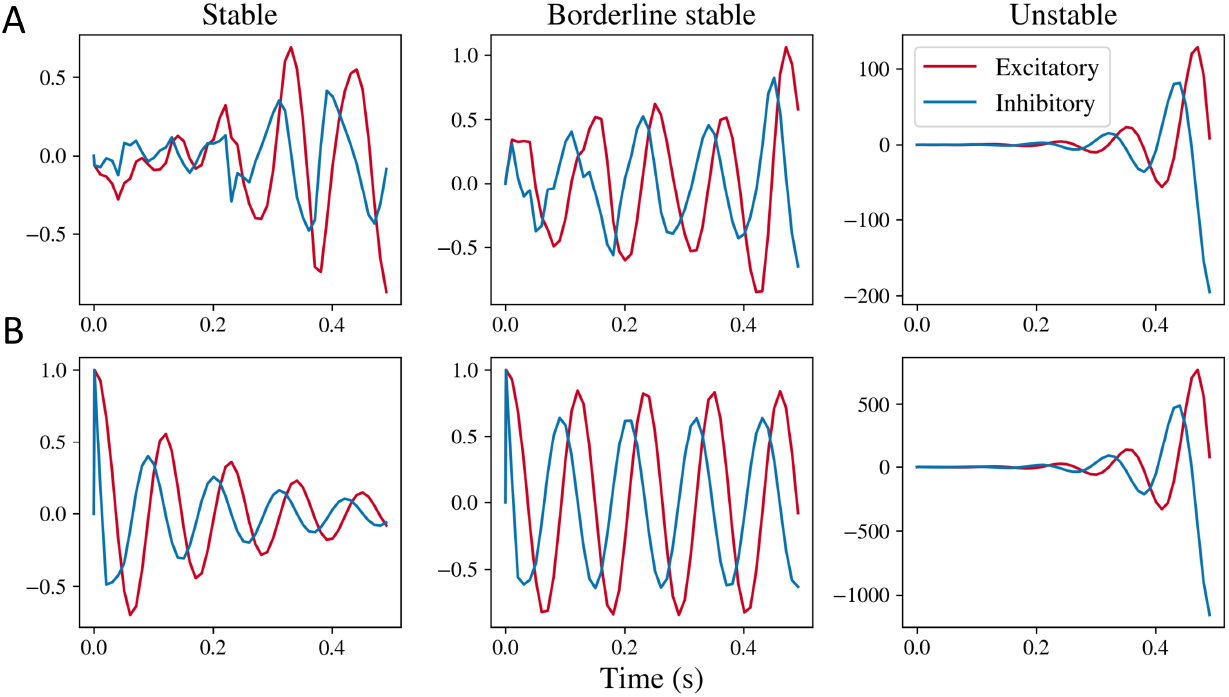
Simulations obtained by taking inverse Laplace transform of the mesoscopic model. **A:** Simulations obtained for the stable, borderline stable approaching limit cycle, and unstable regimes for the complete mesoscopic model including noise. **B:** Simulations obtained for the mesoscopic model’s transfer function with impulse input. Stable simulations are obtained for parameter values *g*_ii_ = 0.5, *g*_ei_ = 0.4, *τ*_e_ = 0.012, *τ*_i_ = 0.003. Borderline stable simulations were obtained for parameter values *g*_ei_ = 0.52, and same values for *g*_i_, *τ*_e_ and *τ*_i_. Unstable simulations were obtained for parameter values *g*_ei_ = 1.0, and same values for *g*_ii_, *τ*_e_, and *τ*_i_.

We generated the time simulations by taking an inverse Laplace of the complete macroscopic model as well. Fig. 2 shows the complete model with noise in (A), frequency response of the complete model without noise but an impulse input in (B), and macroscopic frequency response alone with impulse input in (C). As seen in Fig. 2C, depending on the parameter values, four types of solutions are possible. If *α* > 1, the mean value of the frequency response keeps increasing with time. For certain combinations of *τ*_G_ and *α*, we observe damped oscillations, oscillations that are blowing up with time, and limit cycles. These regimes are clearly distinguishable when simulating the macroscopic impulse response alone. When we include the local model input without noise but ensuring that the local model is stable, stability of the complete model is not obvious. Moreover, the regimes are further unclear when noise is included in the model. We have only demonstrated simulations for initial time points up till around 0.3 seconds because of numerical instabilities encountered while performing inverse Laplace transform for higher time points, which we discuss in the discussions section.

**Figure 2:**
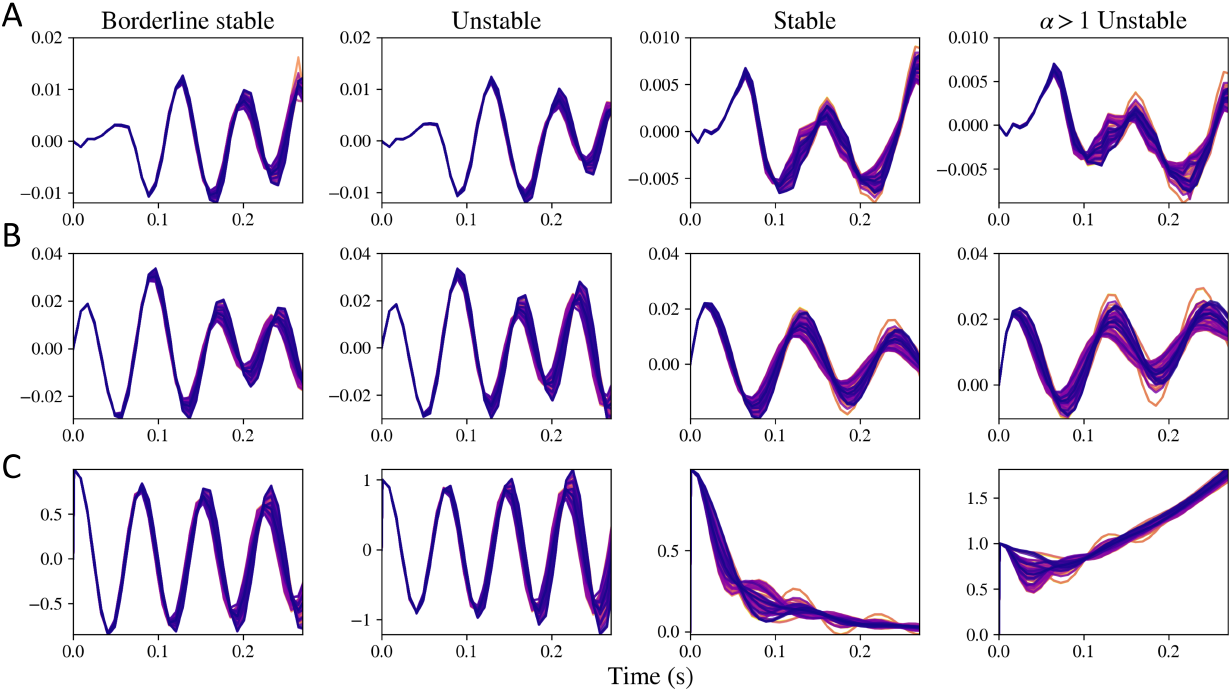
Simulations obtained by taking inverse Laplace transform of macroscopic model. First column shows simulations when the system is nearing the limit cycle and is stable (*τ*_G_ = 0.0055, *α* = 0.1), second column demonstrates system that is unstable because of a low value of *τ*_G_ (*τ*_G_ = 0.005, *α* = 0.1), third columns shows a system that is stable (*τ*_G_ = 0.012, *α* = 0.8), and fourth columns shows a system that is unstable due to a high value of *α* (*τ*_G_ = 0.012, *α* = 1.1). **A:** Complete simulations. **B:** Macroscopic model with the mesoscopic model but without noise and with an impulse input. **C:** Impulse transfer function of the macroscopic model.

### 2.2 SGM generates stable damped oscillatory solutions for a range of parameters

Based on the stability method described, we obtained the stability regimes for the macroscopic as well as the local mesoscopic model alone. The system is stable when the oscillations dampen over time, is a limit cycle if the amplitude of oscillations remain constant over time, and is unstable if the amplitude or mean of the oscillations increase with time. For the mesoscopic model, the stability is largely controlled by the neural gain parameters *g*_ei_ and *g*_ii_, as shown in Fig. 3B. For higher values of *g*_ei_ and *g*_ii_, the system becomes unstable and the oscillations amplitude increase with time as shown in the rightmost column in Fig. 1. The time constants *τ*_e_ and *τ*_i_ only shift the stability boundary marginally. This is because a change in parameters *g*_ei_ or *g*_ii_ can shift the poles of the characteristic polynomial in Eq. (28) to the right of the imaginary axis, as shown in the Fig. 3A. Since the poles that cross the imaginary axis have a non-zero imaginary component as well, the system will exhibit oscillations that blow up with time. As seen in the supplementary Fig. S3B as well, shifting the time constants does not shift the stability boundary substantially. At the stability boundary, we observe the limit cycles type solutions as demonstrated in the middle column in Fig. 1.

**Figure 3:**
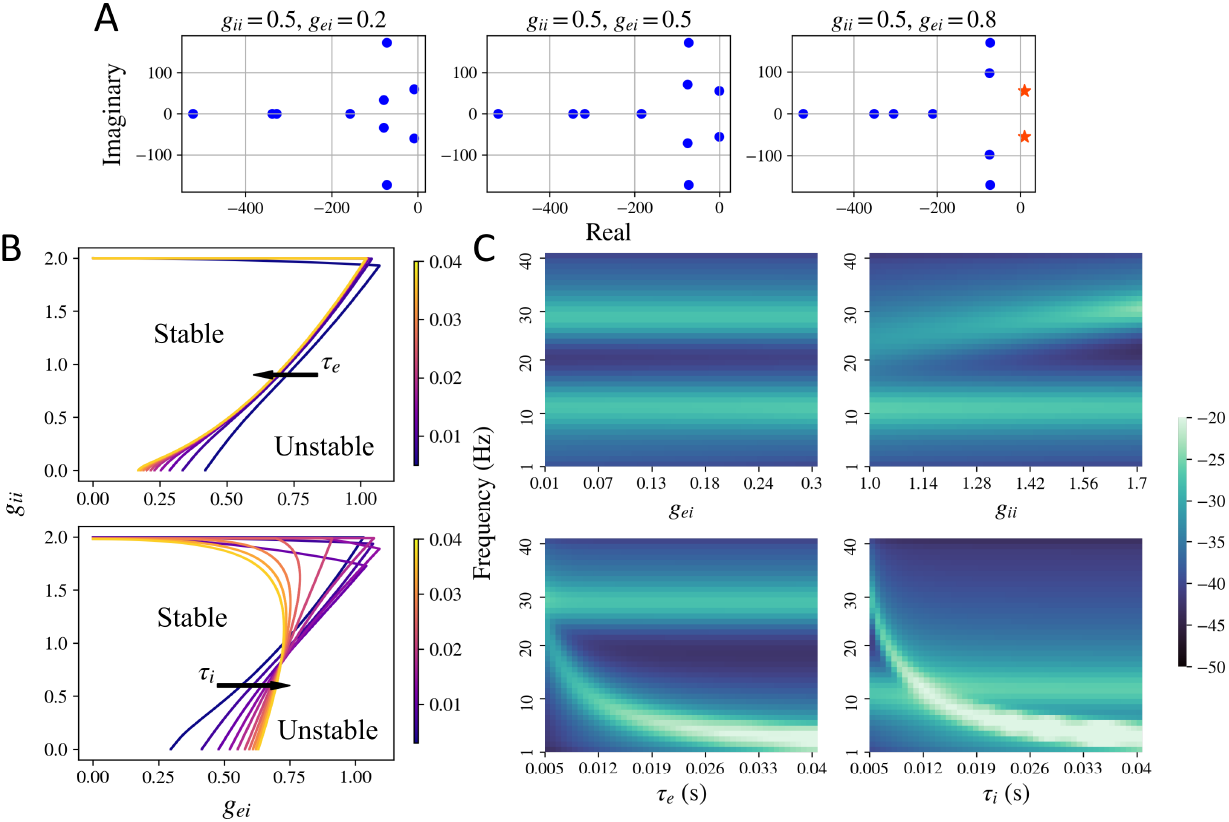
Stability regime for the mesoscopic model. **A:** Plot of the poles of the characteristic equation Eq. (28) for stable and unstable systems. All poles to the left of the imaginary axis are plotted as blue dots and the ones to the right of the imaginary axis are plotted as orange stars. Upon increasing *g*_ei_, a pair of poles shift to the right of the imaginary axis and the system becomes unstable. The poles were generated for default parameters *τ*_e_ = 0.012 and *τ*_i_ = 0.003 s. **B:** Limit cycle boundary obtained for the mesoscopic model for different values of *g*_ei_, *g*_ii_, *τ*_e_, and *τ*_i_. The mesoscopic model is stable for lower values of *g*_ei_ and *g*_ii_. **C:** Frequency spectra observed in the stable regime while varying one of the model parameters and keeping others fixed. Default parameters were set at *g*_ei_ = 0.25, *g*_ii_ = 1.5, *τ*_e_ = 0.01, and *τ*_i_ = 0.005.

The mesoscopic model can also exhibit a variety of peak frequencies, demonstrated in Fig. 3C and supplementary Fig. S1. As shown in Fig. 3C, a primary and a secondary peak in the frequency spectra can be observed, depending on the parameter values. For fixed *τ*_e_ and *τ*_i_ and varying *g*_ei_ and *g*_ii_, a primary alpha and a secondary higher beta peak can be observed. For higher values of *τ*_e_ and *τ*_i_, a primary delta peak can be observed. The local model’s spectra can also exhibit a primary peak in the gamma band, for very low values of *τ*e and *τ*i, as shown in the supplementary Fig. S1.

The stability regime of the macroscopic model is demonstrated in Fig. 4. For the macroscopic model, all the parameters *τ*_e_, *τ*_G_, *α*, and *v* impact the stability. Time constant *τ*_e_ determines the boundary for stability of oscillations when *α* = 0, as demonstrated in the Methods section. Speed *v* determines the shape of the boundary for stability of oscillations which is shown with a blue line. If speed becomes zero, the effect due to the connectivity matrix becomes zero, and the situation is the same as that for *α* = 0. In such a case, the boundary of stability will be a horizontal line at 2*τ*_g_ = *τ*_e_. There is also a hard boundary of stability at *α* = 1 shown as a red line, as was demonstrated in the Methods section. For *α* > 1, the mean of oscillations starts increasing with time, making the system unstable. Moreover, for sufficiently low values of *τ*_g_ and *α* > 1, both the mean and the amplitude of oscillations increase with time. These example regimes are labeled in Fig. 4A and the impulse response transfer function is simulated in Fig. 4B. As seen in the simulations shown in Fig. 4C 1, since *τ*_G_ is sufficiently high and *α* ≤ 1, the oscillations dampen over time. In Fig. 4B 2, since *τ*_G_ is below the stability boundary in blue but *α* ≤ 1, the amplitude of the oscillations increase over time even though the mean of the oscillations does not change. In Fig. 4B 3, since *α* > 1 even though *τ*_G_ is sufficiently high, the mean of the oscillations increase with time even though the oscillations dampen over time. In Fig. 4B 4, since *τ*_G_ is sufficiently low and *α* > 1, both the amplitude and mean of oscillations will increase with time.

**Figure 4:**
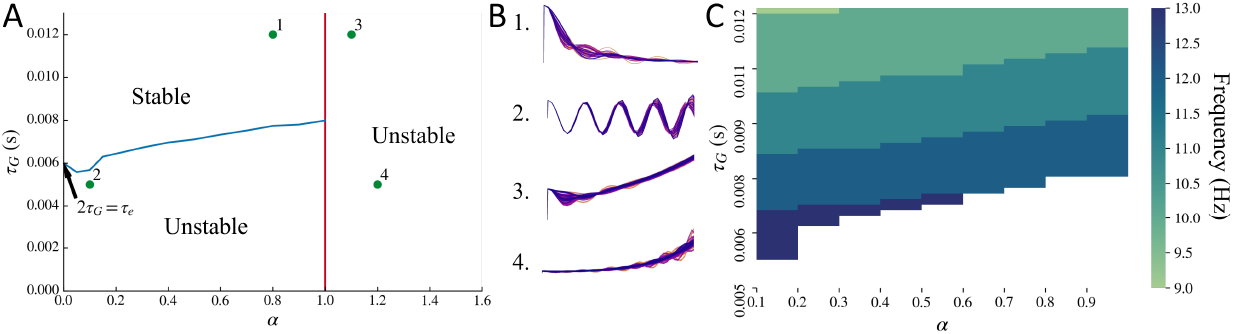
Stability regime for the macroscopic model. **A:** Stability boundary obtained for the macroscopic model. System is unstable for lower values of *τ*_G_ when *α* ≤ 1, shown as the region below by the blue line. System is also unstable for *α* > 1, shown as the region to the right of the red line. The blue line stability boundary was obtained for default parameters *τ*_e_ = 0.012 s and *v* = 5 m/s. **B:** The points marked in A as 1, 2, 3, and 4 are demonstrated in B as time simulations. Note: The simulation plots in B have been stretched out to clearly demonstrate the mean and amplitude of oscillations. **C:** Frequency at which the primary peak is observed in the modeled macroscopic frequency spectra upon varying *τ*_G_ and *α* simultaneously, while replacing mesoscopic model input with exp (−*t*). Parameters *τ*_e_ = 0.012 s and *v* = 5 m/s as default. White region corresponds to the unstable regime. For most of the combinations of *τ*_G_ and *α* for which the system is stable, a peak in the alpha frequency band is observed.

### 2.3 Macroscopic model alone can exhibit a peak in the alpha frequency band

We observed that the macroscopic model’s spectra can exhibit a single peak in the alpha frequency band even without the local mesoscopic oscillatory signals *x*_e_(*t*) + *x*_i_(*t*) as the input to the macroscopic model. To test this, we replaced the local mesoscopic model input of *x*_e_(*t*) + *x*_i_(*t*) with exp (−*t*). This is a simple damping term and its Fourier transform will be 1/(*jω* + 1). Thus, if a peak in a frequency band is observed in this model’s spectra, it can only get generated from the macroscopic response. A peak in the alpha frequency band can be seen for a combination of parameters *τ*_G_ and *α* in the stable regime, as shown in Fig. 4C. Note that this peak is dependent on *τ*_e_, and altering *τ*_e_ will shift the peak frequency.

The macroscopic model can also exhibit a variety of frequencies at which a peak in the spectra is observed, as shown in the supplementary frequency heat maps for different model parameters in supplementary Fig. S2. The primary peak in the macroscopic model can also shift upon varying the time constants *τ*_e_ and *τ*_G_. Note that a secondary peak is not observed in this macroscopic model alone. The secondary peak is exhibited by the local model, shown in Fig. 3C, and therefore is exhibited at the macroscopic level when the local model is included as an input to the macroscopic model.

### 2.4 Dynamics in functional activity is captured by fluctuations of a small set of parameters

Next, we used the time-frequency decomposition of MEG source-reconstructed time series to estimate model parameters over time, approximately every 5 seconds. We only varied *α*, *g*_ei_ and *g*_ii_. The dynamic model parameters are shown in Fig. 5A, B, and C. All the three parameters vary over time for many subjects. Interestingly, a sharp switch in a can be seen for many subjects. To capture this variation, we counted the number of times the difference between two consecutive values of *α* was greater than 0.5 over time for every subject. This is shown in the supplementary Fig. S4. For multiple subjects, sharp switches occur. Parameter *α* captures the extend of global connectivity.

**Figure 5:**
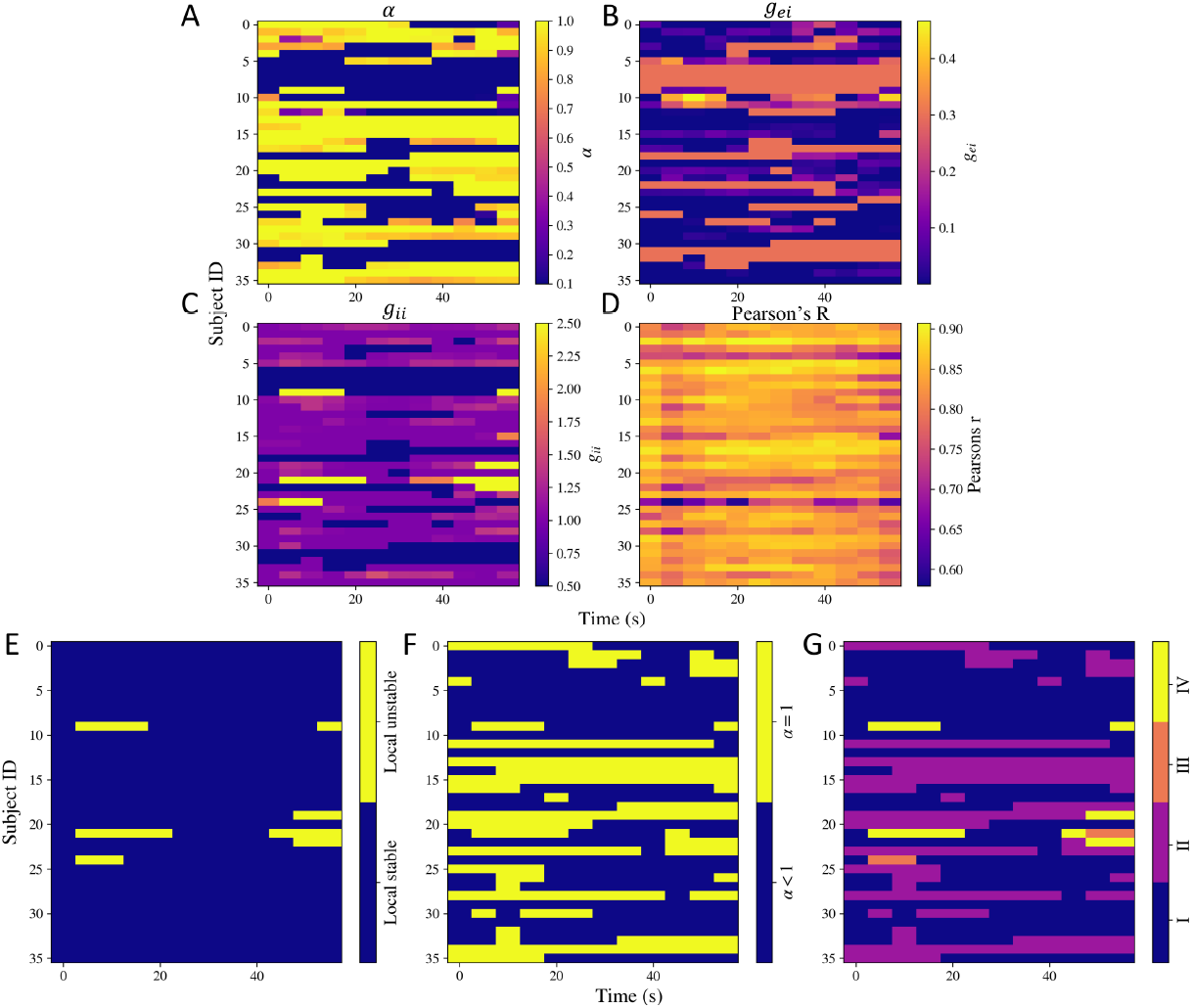
Dynamic model parameters. **A:** *α*, **B:** *g*_ei_, **C:** *g*_ii_, and **D:** the goodness of fit Pearson’s R calculated at different time points for all the subjects. **E, F, G: Dynamic stability. E:** Stability of the local model over time. **F:** Switches in *α* over time. **G:** Switches in different regimes of stability over time. The shade is based on 4 situations: I) Both local model is stable and *α* < 1, II) local model is stable but *α* = 1, III) local model is unstable but *α* < 1, IV) local model is unstable and *aα* = 1.

For the previous model parameter estimation, we kept an upper bound of 1.0 on *α*, to ensure stability. We also estimated how the model parameters vary if the upper bound is relaxed. In supplementary Fig. S5A, B, and C, we show how the model parameters vary when the upper limit of *α* has been increased to 3. We again see switches in *α*, which we show in supplementary Fig. S6.

We also tested if having static parameters instead will be equally accurate in capturing the dynamic spectra. For this, we calculated the Pearson’s correlation coefficient between the modeled spectra with static parameters and the spectra at each of the time points. Then, we performed a paired t-test between the two set of correlations after performing a Fisher’s z transform on them. Based on a one-sided paired t-test, having dynamic parameters gave significantly higher Pearsons’r as compared to having static model parameters, with ap value *p* < 0.0001.

### 2.5 Dynamics in functional activity is captured by fluctuations in model stability

Based on the estimated dynamic model parameters, we also calculated the stability at the respective time points. Note that the neural gain terms control the local model’s stability, while *α* controls the macroscopic model’s stability. By keeping an upper bound of 1 on *α*, we ensured the macroscopic system does not become unstable because of increase in α. The dynamic stability patterns are shown in Fig. 5E, F, and F. As shown in Fig. 5E, the local model’s stability varies with time for very few subjects. As shown in Fig. 5F, *α* hits the stability boundary of *α* = 1 over time for some subjects too. In order to capture if the local and macroscopic model’s stability was varying simultaneously, we show their combination in Fig. 5G. The shade is based on 4 situations: I) Both local model is stable and *α* < 1, II) local model is stable but *α* = 1, III) local model is unstable but *α* < 1, IV) local model is unstable and *α* = 1. This plot shows that most changes in stability patterns occur because of *α* hitting the upper bound.

We repeated this analysis with an increased upper bound of *α* = 3. This is shown in supplementary Fig. S5E, F, and G. Similar changes in the dynamics are observed here as well. In particular, *α* switches between stable and unstable regimes for many subjects. As a consequence, we see that the macroscopic model goes unstable while the local model remains stable at different time points, shown as the pink region in Fig. S5G.

We note that the values of *τ*_e_ and *τ*_G_ also control the macroscopic model’s stability, which are static. It implies either the system is constantly stable or unstable depending on their values. They are shown in supplementary Fig. S7. For all the subjects except two, the macroscopic system is stable. Constraining the time constants appropriately is beyond the scope of this work, but we will investigate it in the future.

## 3 Discussion

In this work, we demonstrated that a biophysical linearized spectral graph model can generate frequency-rich spectra. The key advantage of this model is that it is hierarchical, analytic, graph-based, and consist of a parsimonious set of biophysically interpretable global parameters. Although prior work has already demonstrated the SGM’s ability to fit wideband regional power spectra, its ability to accommodate more complex dynamics including regimes of stability and instability were previously unknown. In this paper we focused on those aspects, along with dynamic changes in model parameters that may then lead to dynamic complex behavior. Using detailed analytical and numerical analyses, we were able to show that this model can exhibit oscillatory solutions that are damped, limit cycles, or unstable, which we demonstrated by calculating the inverse Laplace transform of the model responses. We also showed how the stability of both the mesoscopic (local circuits) and the macroscopic (whole brain network) model are governed by the model parameters. Interestingly, the macroscopic model alone can exhibit a peak in the alpha frequency band even when the local model is replaced with a simple damping term, implying that the macroscopic alpha rhythm may not arise form local oscillators tuned to the alpha frequency, but emerge from the modulatory effect of long-range network connectivity. In addition, a variety of frequency responses can be observed by varying the model parameters within physiological ranges, making this model suitable for inferring model parameters using MEG wideband spectra directly, instead of using second-order metrics such as functional connectivity as previously employed in various non-linear modeling approaches. This will be specifically helpful in capturing differences in frequency spectra observed in different diseases and brain states. Lastly, we inferred dynamic model parameters using time-frequency decomposition of the source-reconstructed MEG data, outlining a novel model-based approach to directly infer dynamics in functional activity using a parsimonious set of biophysically interpretable model parameters.

### 3.1 Relationship to previous works

All the structure-function models can be categorized into communication and control models, reviewed in detail by Srivastava et al. [52]. The dynamical communication models incorporate biophysics of signal propagation and generation. While such models can be linear as well as non-linear, most structure-function modeling approaches are based on non-linear models – such models can exhibit a rich dynamical repertoire in their oscillatory behavior [26–28]. Such behaviors are quantified in terms of bifurcations defining solution regimes that are fixed points, limit cycles, quasiperiodic, chaotic, or bistable. These models have been used extensively and applied to differentiate different brain states and allow transitions between them [53]. While SGM cannot exhibit such complex behavior, it can accurately capture the wideband frequency spectra, in contrast to the other modeling approaches that infer model parameters using second order statistics such as functional connectivity. It is also to be noted that even if the non-linear models can exhibit diverse solutions, they may not be completely derived from biophysics, e.g., some models are based on using normal form of Hopf bifurcation model to represent the mesoscopic dynamics [54, 55].

On the other hand, the controls models are primarily linear time-invariant (LTI) systems. Control models have also been widely used to model state transitions and different neurological conditions with a view of estimating controllability in terms of the energy required to facilitate state transitions [31–34]. Such models are limited in the kinds of solutions they can exhibit – exponential growth, exponential decay, and sinusoidal oscillations. Moreover, these solutions are primarily based on eigenvalues of the structural connectome matrix. While SGM can also exhibit broadly the same kind of solutions that the control models can, it can generate wideband frequency spectra that can accurately match empirical MEG spectra for a range of model parameters. Moreover, SGM is derived from the excitatory and inhibitory neuronal biophysics, unlike the network control models. Thus, SGM can be interpreted as an LTI network control system where the structural connectome is replaced by an eigendecomposition of the complex Laplacian along with the macroscopic excitatory frequency response, and the input control is replaced by the mesoscopic excitatory and inhibitory dynamics.

Numerous evidences have pointed towards the brain being multistable [56] and several modeling approaches have indicated that the brain functional activity exhibits multistability by operating close to a bifurcation point [57, 58, 26, 59, 54]. In such cases, input noise can shift the non-linear solution regime if it is close to a bifurcation point which can yield the dynamical repertoire of simulated functional activity [60, 58, 26, 59]. SGM cannot exhibit multistability in its current form – noise will simply act as a linear filter which shapes the power spectrum. Instead, we focus on capturing fluctuations in functional activity by inferring SGM model parameters at different time points – an alternative approach of introducing non-linearity while keeping parameter inference tractable and ensuring estimation of wideband frequency spectra instead of second order statistics such as functional connectivity.

A key question then arises is that if SGM is sufficient to capture brain macroscopic dynamics without multistability and other complex dynamics. Our modeling approach is focused on capturing the macroscopic spatial and frequency patterns, which can be largely identical across individuals [61–63]. It has been suggested that emergent long-range activity can be independent of microscopic local activity of individual neurons [64, 61, 23, 9, 65, 19], and that these long-range activities may be regulated by the long-range connectivity [66–69]. Therefore, to capture such phenomena, it may be sufficient to undertake deterministic modeling approaches such as SGM. Indeed, it was already demonstrated that SGM outperforms a Wilson-Cowan neural mass model in fitting the empirical MEG spectra [35]. In addition, a recent comparison showed that linear models outperformed non-linear models in predicting resting state fMRI time series. This was attributed to the linearizing affects of macroscopic neurodynamics and neuroimaging due to spatial and temporal averaging, observation noise, and high dimensionality [70]. These evidences, in addition to our results on the dynamic model parameter estimation, suggest that it can be sufficient to use SGM to capture static as well as temporal fluctuations in the functional activity.

Previous MEG studies have reported MEG fluctuations in the order of seconds [38, 71] as well as a much lower order of 100 ms [41]. Presented model parameter variability results were resolved at 5 sec due to the chosen window length. Dynamics on faster timescales will require finer Morlet wavelet time-frequency decomposition. New approaches that can detect non-evenly-spaced state switching [44, 41] may also be adapted in future iterations of our inference procedure.

### 3.2 Potential applications and future work

This work can be extended to identify temporal state and stability changes in the functional activity of various brain states and diseases, particularly neuropsychiatric disorders. We will also examine the association of switching in *α* coupling, which controls the coupling between remote populations, with the dynamics of segregation versus integration. Currently, in our model macroscopic stability is ensured via an upper bound on the coupling term *α*; in the future we will explicitly introduce automatic gain control for this purpose, e.g. as a mathematical correlate of neuromodulation [72].

The presented model-based inference of dynamic functional activity can provide additional insights into the biophysics because of the parsimony and the biophysical interpretability of the SGM parameters. For example, we recently applied the mesoscopic model to estimate regionally varying local model parameters for empirical static MEG spectra collected for healthy and Alzheimer’s disease subjects. Our results showed that the neural gains and the time constants were differentially distributed in the healthy versus the Alzheimer’s disease subjects, indicating an excitatory/inhibitory imbalance in Alzheimer’s disease (*submitted*). Lastly, one can also extend this work to find seizure onset regions, based on the stability of the regional model parameters, as has been done earlier by investigating bifurcation points with non-linear modeling [73].

### 3.3 Limitations

The SGM model involves various limitations and assumptions that have previously been described. Here we list potential limitations of the current stability results. We employed an inverse Laplace transform approach to generate model solution in the time domain, instead of solving the differential equations in the time domain directly. While this is an excellent feature enabled by having a closed-form solution in Laplace domain, a limitation of this approach is that the inverse Laplace transform can lead to numerical instabilities. Hence we were able to obtain reliable solutions only for a short range of time. This is especially true when the solution is close to the limit cycle, exhibiting persistent oscillations, since such systems are harder to invert using numerical inverse Laplace [74]. However, the time range used in this study (0 – 0.3 seconds) is sufficient to demonstrate the nature of the oscillations and whether they will blow up or damp down with time.

We obtained the oscillation stability boundary (blue line in Fig. 4A) as a function of *τ*_G_ and *α*, keeping all other parameters fixed. This boundary will shift upon varying parameters *τ*_e_ and *v* as well, which we have not demonstrated here. Moreover, this is only an approximate boundary since this was based on a numerical root finding approach. This requires an initial guess close to the actual root. It is computationally infeasible currently to perform a symbolic manipulation of an 86X86 matrix in Eq. (37), and to either apply Routh-Hurwitz criteria or find roots of the determinant of this matrix. With sufficient computation capability in the future, an accurate stability boundary may be obtained via the characteristic polynomial of Eq. (37). Parameter estimation of two subjects gave *τ*_G_ ≪ *τ*_e_, indicating that their inferred macroscopic system is unstable. In future work we will explore constraints to ensure parameters are in the stable regime.

In this study, we only considered 1 minute MEG recordings; the quality of inferred dynamics might therefore benefit from longer recordings. However, we chose the most noise-free 1 minute snippet out of the entire 5 minute recording, ensuring that the fluctuations observed in model parameters are not because of measurement noise.

## 4 Theory and Methods

### Notation

All the vectors and matrices are written in boldface and the scalars are written in normal font. The frequency *f* of a signal is specified in Hertz and the corresponding angular frequency *ω* = 2*πf* is used to obtain the Fourier transforms. The connectivity matrix is defined as ***C*** = *c_jk_*, where *c_jk_* is the connectivity strength between regions *j* and *k,* normalized by the row degree.

### 4.0.1 Mesoscopic model

For every region *k* out of the total *N* regions, we model the local excitatory signal *x_e,k_*, local inhibitory signal *x*_i,*k*_ as well as the long range excitatory signal *x_k_* where the global connections are incorporated. The local signals are modeled using an analytical and linearized form of neural mass equations. We write a set of differential equations for evolution of *x_e,k_* and *x*_i,*k*_ due to decay of individual signals with a fixed neural gain, incoming signals from populations that alternate between excitatory and inhibitory signals, and input white Gaussian noise. Letting *f_e_*(*t*) and *f_i_*(*t*) denote the ensemble average neural impulse response functions, the *x*_e,*k*_ and *X*_i,*k*_ are modeled as

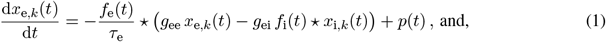

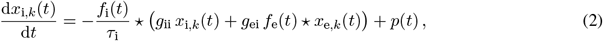

where, ⋆ stands for convolution, *p*(*t*) is input noise, parameters *g*_ee_, *g*_ii_, *g*_ei_ are neural gain terms, and parameters *τ*_e_, *τ*_i_ are characteristic time constants. These are global parameters and are the same for every region *k*. Here, the ensemble average neural impulse response functions *f_e_*(*t*) and *f*_i_(*t*) are assumed to be Gamma-shaped and written as

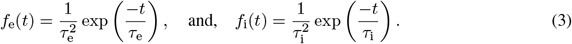

### 4.0.2 Macroscopic model

A similar equation is written for the macroscopic signal *x_k_*, for every *k^th^* region, accounting for long-range excitatory corticocortical connections for the pyramidal cells. The evolution of *x_k_* is assumed as a sum of decay due to individual signals with a fixed excitatory neural gain, incoming signals from all other connected regions determined by the white matter connections, and the input signal *x_e,k_*(*t*) + *x*_i,*k*_(*t*) determined from Eq. (1), (2). Signal *x_k_* is modeled as

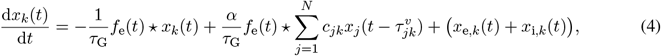

where, *τ*_G_ is the graph characteristic time constant, *α* is the global coupling constant, *c_jk_* are elements of the connectivity matrix determined from DTI followed by tractography, 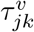 is the delay in signals reaching from the *j^th^* to the *k^th^* region, *v* is the cortico-cortical fiber conduction speed with which the signals are transmitted. The delay 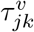 is calculated as *d_jk_*/*v*, where *d_jk_* is the distance between regions *j* and *k*.

These set of equations are parameterized by 8 global parameters: excitatory time constant *τ*_e_, inhibitory time constant *τ*_i_, macroscopic graph time constant *τ*_G_, excitatory neural gain *g*_ee_, inhibitory neural gain *g*_ii_, alternating population neural gain *g*_ei_, global coupling constant *α*, and conduction speed *v*. The neural gain *g*_ee_ is kept as 1 to ensure parameter identifiability. We estimate the 7 global parameters using an optimization procedure described in the next section.

### 4.0.3 Model solution in the Fourier domain

Since the above equations are linear, we can obtain a closed-form solution in the Fourier domain as demonstrated below. The Fourier transform 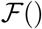 is taken at angular frequency *ω* which is equal to 2*πf*, where *f* is the frequency in Hertz. The Fourier transform of the equations (1), (2) will be the following, where 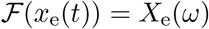 and 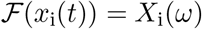, and j is the imaginary unit:

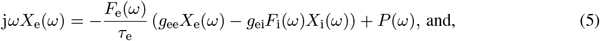

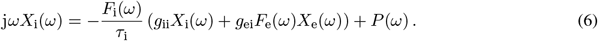

Here, *P*(*ω*) is the Fourier transform of the input Gaussian noise *p*(*t*) which we assume to be identically distributed for both the excitatory and inhibitory local populations for each region, and the Fourier transforms of the ensemble average neural response functions are

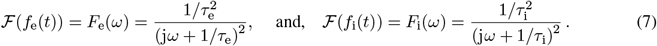

On solving the above equations (5) and (6), *X*_e_(*ω*) and *X*_i_(*ω*) are

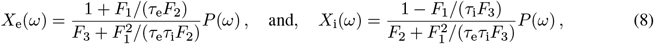

where,

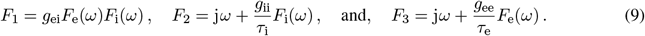

Then, the transfer functions *H*_e_(*ω*) and *H*_i_(*ω*) can be separated out and we get

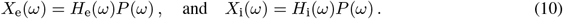

The total neural population is therefore

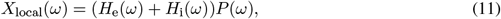

thus, *H*_local_(*ω*) = *H*_e_(*ω*) + *H*_i_(*ω*).

In order to obtain a Fourier response of the macroscopic signal, we first re-write Eq. (4) in the vector form

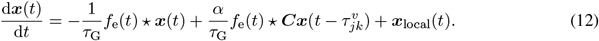

We similarly take a Fourier response of the macroscopic signal and obtain the following as the Fourier transform of Eq. (12), where 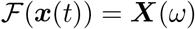:

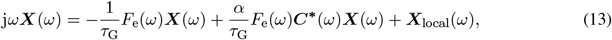

where, 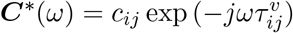. Note that each element in the matrix ***C*** is normalized already by the row degree. The above equation can be re-arranged to give

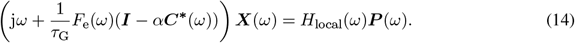

Here, we define the complex Laplacian matrix:

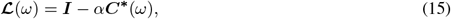

where, *I* is the identity matrix of size *N* × *N*. The eigen-decomposition of this complex Laplacian matrix is

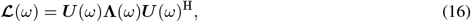

where, ***U***(*ω*) are the eigenmodes/eigenvectors and **Λ**(*ω*) = diag([λ_1_(*ω*),…, λ_*N*_(*ω*)]) consist of the eigenvalues λ_1_(*ω*),…, λ_*N*_(*ω*), at angular frequency *ω*. The macroscopic response Eq. (14) becomes

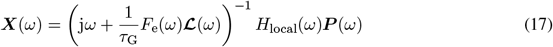

By using the eigen-decomposition of the Laplacian matrix, this yields

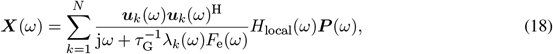

where, ***u***_*k*_(*ω*) are the eigenmodes from **U**(*ω*) and λ_*k*_(*ω*) are the eigenvalues from **Λ**(*ω*) obtained by the eigen-decomposition of the complex Laplacian matrix 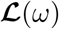 obtained in Eq. 16. Equation (18) is the closed-form steady state solution of the macroscopic signals at a specific angular frequency *ω*. We use this modeled spectra to compare against empirical MEG spectra and subsequently estimate model parameters.

### 4.1 Model parameter estimation

The dataset used for this work is based on the preprocessed publicly available dataset for the SGM work [75], and is also the same as the one we used for the modified SGM [49]. For this dataset, MEG, anatomical MRI, and diffusion MRI was collected for 36 healthy adult subjects (23 males, 13 females; 26 left-handed, 10 right-handed; mean age 21.75 years, age range 7-51 years). Data collection procedure has already been described previously [35]. All study procedures were approved by the institutional review board at the University of California at San Francisco and are in accordance with the ethics standards of the Helsinki Declaration of 1975 as revised in 2008. MEG recordings were collected for 5 minutes while the subjects were resting and had eyes closed. Out of the 5 minute recording, a 1 minute snippet was chosen which was most noise free. MRI followed by tractography was used to generate the connectivity and distance matrices. The publicly available dataset consisted of processed connectivity and distance matrices, and power spectral density (PSD) for every subject. MEG recordings were downsampled to 600 Hz, followed by a bandpass filtering of the signals between 2 to 45 Hz using `firls` in MATLAB [76] and generation of the static frequency spectra for every region of interest using the pmtm algorithm in MATLAB [76]. For generating the the time-frequency decomposition of the MEG time series, Morlet wavelet algorithm in python was used with the input parameter w as 600 and the widths were calculated based on w for every frequency between 2 and 45 Hz.

Modeled spectra was converted into PSD by calculating the norm of the frequency response and converting it to dB scale by taking 20log_10_() of the norm. Pearson’s correlation coefficient between modeled PSD and the MEG PSD was used a goodness of fit metric for estimating model parameters. Pearson’s correlation coefficient was calculated for comparing spectra between each of the regions, and then the correlation coefficient was averaged over all the 68 cortical regions. This average correlation coefficient was the objective function for optimization and used for estimating the model parameters. We used a dual annealing optimization procedure in Python and performed parameter optimization [77]. Parameter initial guesses and bounds for estimating the static spectra are specified in Table 1. Dynamic model parameters were estimated approximately every 5 seconds by fitting the modeled spectra to the frequency spectra every 5 seconds from the time-frequency decomposition. Parameter initial guesses and bounds for estimating the dynamic spectra are specified in Table 2. In this case, the time constants and speed were kept the same as those estimated from the static spectra for every subject. Only *α*, *g_ei_*, and *g_ii_* were allowed to vary. The dual annealing optimization was performed for three different initial guesses, and the parameter set leading to maximum Pearson’s correlation coefficient was chosen for each subject. The dual annealing settings were: `maxiter = 500`. All the other settings were the same as default.

**Table 1:** SGM parameter values, initial guesses, and bounds for parameter estimation for static spectra fitting

| Name | Symbol | Initial value 1 | Initial value 2 | Initial value 3 | Lower/upper bound for optimization |
| --- | --- | --- | --- | --- | --- |
| Excitatory time constant | $\tau_e$ | 0.012 s | 0.018 s | 0.006 s | [0.005 s, 0.02 s] |
| Inhibitory time constant | $\tau_i$ | 0.003 s | 0.01 s | 0.018 s | [0.005 s, 0.02 s] |
| Long-range connectivity coupling constant | $\alpha$ | 1 | 0.5 | 0.1 | [0.1, 1] |
| Transmission velocity | $v$ | 5 m/s | 10 m/s | 18 m/s | [5 m/s, 20 m/s] |
| Alternating population gain | $g_{ei}$ | 0.2 | 0.1 | 0.3 | [0.001, 0.8] |
| Inhibitory gain | $g_{ii}$ | 1 | 1.5 | 0.5 | [1, 2.5] |
| Graph time constant | $\tau_G$ | 0.006 s | 0.01 s | 0.018 s | [0.005 s, 0.02 s] |
| Excitatory gain | $g_{ee}$ | n/a | n/a | n/a | n/a |

**Table 2:** SGM parameter values, initial guesses, and bounds for parameter estimation for dynamic spectra fitting

| Name | Symbol | Initial value 1 | Initial value 2 | Initial value 3 | Lower/upper bound for optimization |
| --- | --- | --- | --- | --- | --- |
| Long-range connectivity coupling constant | $\alpha$ | 1 | 0.5 | 0.1 | [0.1, 1] |
| Alternating population gain | $g_{ei}$ | 0.2 | 0.1 | 0.3 | [0.001, 0.8] |
| Inhibitory gain | $g_{ii}$ | 1 | 1.5 | 0.5 | [1, 2.5] |

### 4.2 Time Domain simulations

The closed-form solution obtained in Eq. (18) represents the steady-state response for a specific angular frequency *ω*. In order to obtain the transient behavior of the model, we performed an inverse Laplace transform of the model solution obtained in the Laplace domain. The closed-form solution in the Laplace domain is the same as that in Eq. (18), replacing *jω* with *s*, where *s* is the Laplace variable which gives

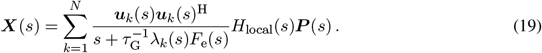

Here, *H*_local_(*s*) = *H*_e_(*s*) + *H*_i_(*s*), and *P*(*s*) is the Laplace transform of the input noise term *p*(*t*).

We performed inverse Laplace transform of the equations using python’s mpmath’s numerical inverse Laplace algorithm, based on the de Hoog, Knight, and Stokes method [78].

### 4.3 Stability analysis of the mesoscopic model

We performed a stability analysis of the model and explored regimes demonstrated in Fig. 1 and 2. Firstly, we explored the stability of the mesoscopic model alone. Then, we investigated the stability of the macroscopic model. For performing the stability analysis, we obtained the set of equations in Laplace domain. Then, we identified the poles on the impulse transfer function, where the poles imply the roots of the denominator of the impulse transfer function. If any of the poles were to the right of the imaginary axis, the system will be unstable.

The local excitatory system in the Laplace domain is

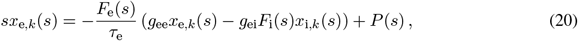

and the local inhibitory system in the Laplace domain is

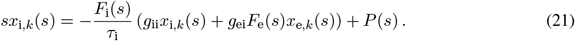

Here,

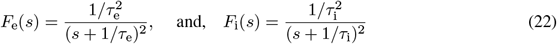

are the neural impulse response functions in the Laplace domain, and *P*(*s*) is the Laplace transform of Gaussian white noise. Equations (20) and (21) can be written in matrix form:

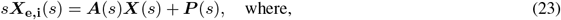

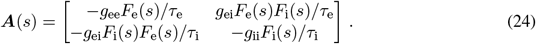

Here, ***X*_e,i_**(*s*) ≡ [*x*_e,*k*_(*s*), *x*_i,*k*_(*s*)]. This yields

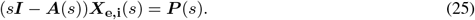

Therefore, the poles of the solution will be the roots of the determinant |*s****I*** – ***A***(*s*)|, which is

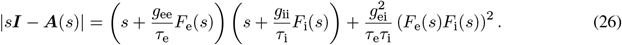

Letting *t*_e_ = 1/*τ*_e_, *t*_i_ = 1/*τ*_i_, we have 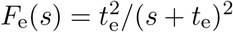 and 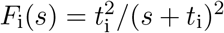, we get

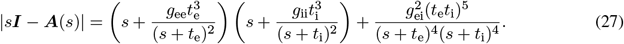

By multiplying the right hand side of the above equation with (*s* + *t*_e_)^4^(*s* + *t*_i_)^4^, we get the following polynomial in *s*:

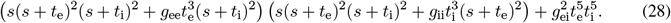

This is the characteristic polynomial of the mesoscopic model. We numerically solve for and find the roots of this polynomial, which are also called the poles. If the real part of any of the roots is positive, it will imply that the system is unstable. Another way to check for stability is the Routh-Hurwitz criterion. If there are any changes in the sign of the elements of the Routh-Hurwitz array, it would imply that the system is unstable. We used both numerical root finding and the Routh-Hurwitz criterion to check for stability. The roots of the polynomial or the poles are plotted in Fig. 3A. As seen in this figure, upon increasing the neural gains, a pair of poles cross the imaginary axis and the real part becomes greater than zero. This loss in stability was also confirmed by calculating the Routh-Hurwitz array.

### 4.4 Stability analysis of the macroscopic model

Since the macroscopic model consists of 86 regions as defined by the Desikan-Killiany atlas [50], it is numerically not feasible to obtain a characteristic polynomial of the determinant and subsequently its roots to investigate stability. Here, we use different approaches to obtain the stability boundary. We investigate two different scenarios: i) *α* = 0, and ii) *α* > 0, separately below.

#### 4.4.1 Stability boundary when *α* = 0

First, we check if the macroscopic model is stable even with *α* = 0. With *α* = 0, the macroscopic model equations will become

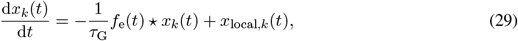

for every region *k*. Its Laplace transform is given by

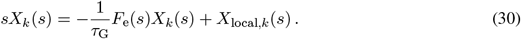

Solving this equation yields

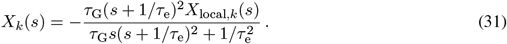

Since we can obtain a characteristic polynomial in this case, we can analytically investigate stability. The characteristic polynomial is

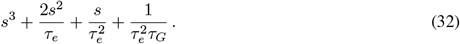

According to the Routh-Hurwitz criteria for a 3-degree polynomial, for stability, 2/*τ*_e_ and 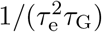 should be positive and the following condition should hold:

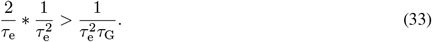

Therefore, the condition for stability when *α* = 0 is

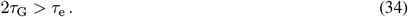

We confirmed this condition by estimating the poles using numerical root finding as well. This provides us a boundary for stability when *α* = 0. Next, we will use both analytical and numerical approaches to estimate stability boundaries when *α* > 0.

#### 4.4.2 Stability boundary when *α* > 0

For the macroscopic model with *α* > 0, the Laplace transformed equation will be

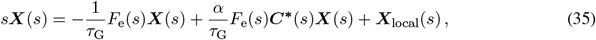

where, 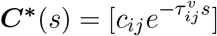. This can be re-written as

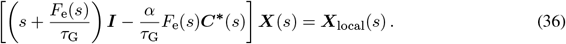

The roots of the determinant of the following matrix determines the stability of the system:

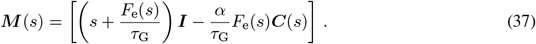

We will find boundaries where at least one of the roots will cross the imaginary axis. To this end, we will investigate two sub-cases separately: i) *s* = 0 is a root of ***M***(*s*), and ii) *s* = *jω* is a root of ***M***(*s*) for some value of *ω*. Both *s* = 0 and *s* = *jω* will define the stability boundaries. We will also use the intuition that the system will ultimately become unstable for high values of *α* since it controls the signal input to the system. Therefore, we only need to find the upper bound on *α* to determine the stability boundary.

##### Case i)

*s* = 0 **is a root of *M***(*s*) We will first check when *s* = 0 is a root of the determinant of ***M***(*s*). When *s* = 0, the determinant becomes

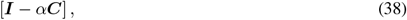

here, ***C*** is a real adjacency matrix normalized by row degree. For this determinant to be zero, one of the eigenvalues λ_*i*_ of [***I*** – *α**C***] should be zero. Since **C** is a normalized adjacency matrix, from graph theory, we know that

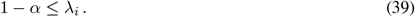

One of the eigenvalues will be zero when *α* = 1. Therefore, *α* = 1 is the stability boundary, and *α* ≥ 1 will make the system unstable, regardless of other parameter values.

##### Case ii)

*s* = *jω* **is a root of *M***(*s*) **for some value of** *ω* In this case, we need to solve the above determinant [***I*** – *α**C***] and find values of *τ*_G_ and *α* for which *s* = *jω* is a root of the determinant, for some value of *ω*. We used numerical root finding, giving different initial guesses for *τ*_G_ and *α*. In particular, we used the hybr method in Python’s Scipy’s root finding library, with these settings: `xtol = 1e-12`, `maxfev = 10000` [79]. Based on this, we generated a stability regime by varying *τ*_G_ and *α*. Note that all other parameters were fixed while finding these stability boundaries. These boundaries will shift when the parameters *τ*_e_ and *v* are varied. However, the upper bound of *α* = 1 will remain unaffected since that was found analytically earlier and it was not based on any other parameter value.

## Data availability

The code and processed datasets for this work can be found in this github repository: https://github.com/Raj-Lab-UCSF/spectrome-stability.

## Acknowledgments

This work was supported by NIH grants R01NS092802/ 183412 and RF1AG062196. The template HCP connectome used in the preparation of this work were obtained from the MGH-USC Human Connectome Project (HCP) database (https://ida.loni.usc.edu/login.jsp). The HCP project is supported by the National Institute of Dental and Craniofacial Research (NIDCR), the National Institute of Mental Health (NIMH) and the National Institute of Neurological Disorders and Stroke (NINDS). Collectively, the HCP is the result of efforts of co-investigators from the University of Southern California, Martinos Center at Massachusetts General Hospital (MGH), Washington University, and the University of Minnesota.

## Supplementary Figures

**Figure S1:**
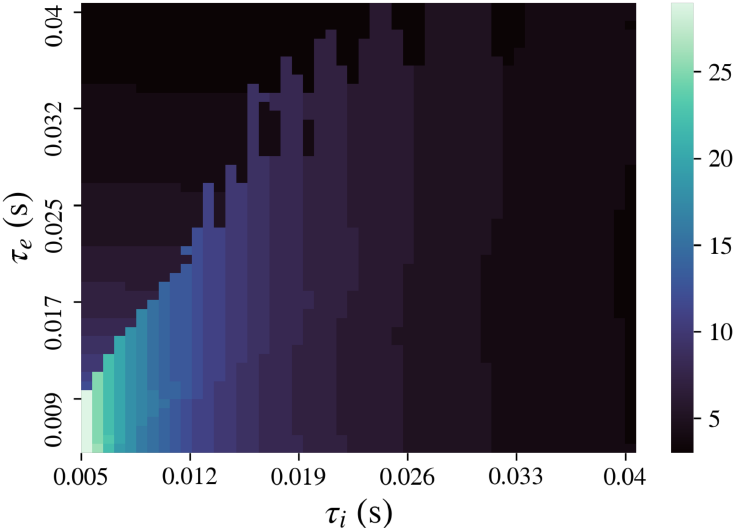
Frequency at which the primary peak is observed in the modeled mesoscopic frequency spectra upon varying *τ*_e_ and *τ*_i_ simultaneously. Peak in the gamma region can be obtained for very low values of *τ*_e_ and *τ*_i_ both. For most of the parameter values, a peak in the lower frequency range is observed. Here, *g*_ii_ = 1.5 and *g*_ei_ = 0.25 to ensure stability.

**Figure S2:**
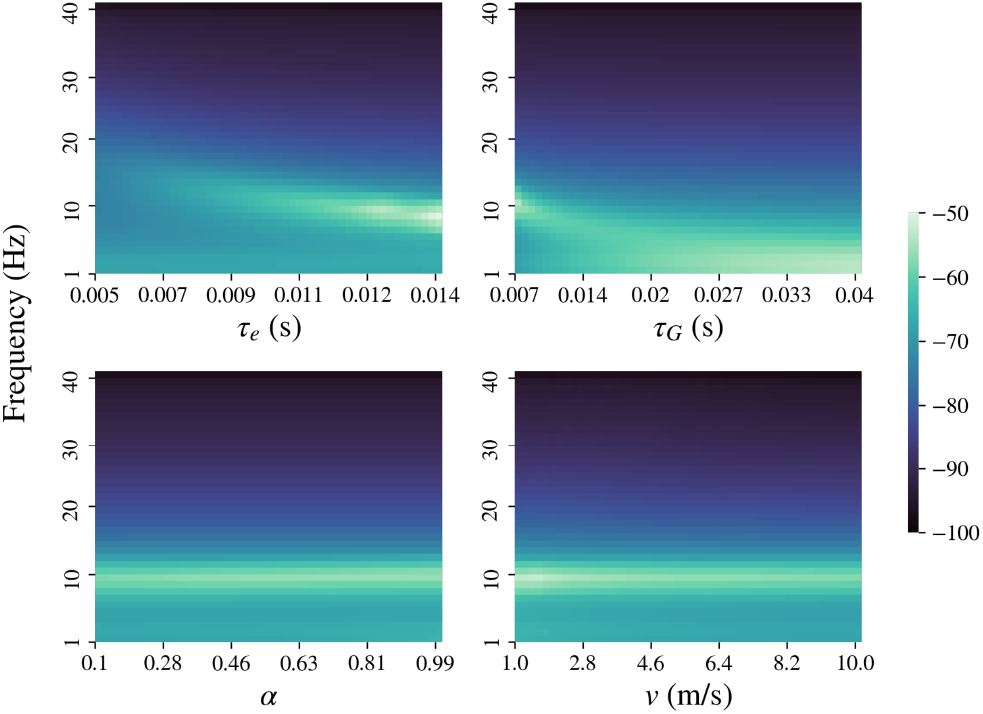
Macroscopic modeled frequency spectra obtained by varying model parameters: **A:** *τ*_e_, **B:** *τ*_G_, **C:** *α*, and **D:** *v*. This spectra is obtained when the local input *x*_e_(*t*) + *x*_i_(*t*) is replaced with exp (−*t*). The default parameters are *α* = 0.5, *v* = 5 m/s, *τ*_e_ = 0.012, and *τ*_G_ = 0.008 s.

**Figure S3:**
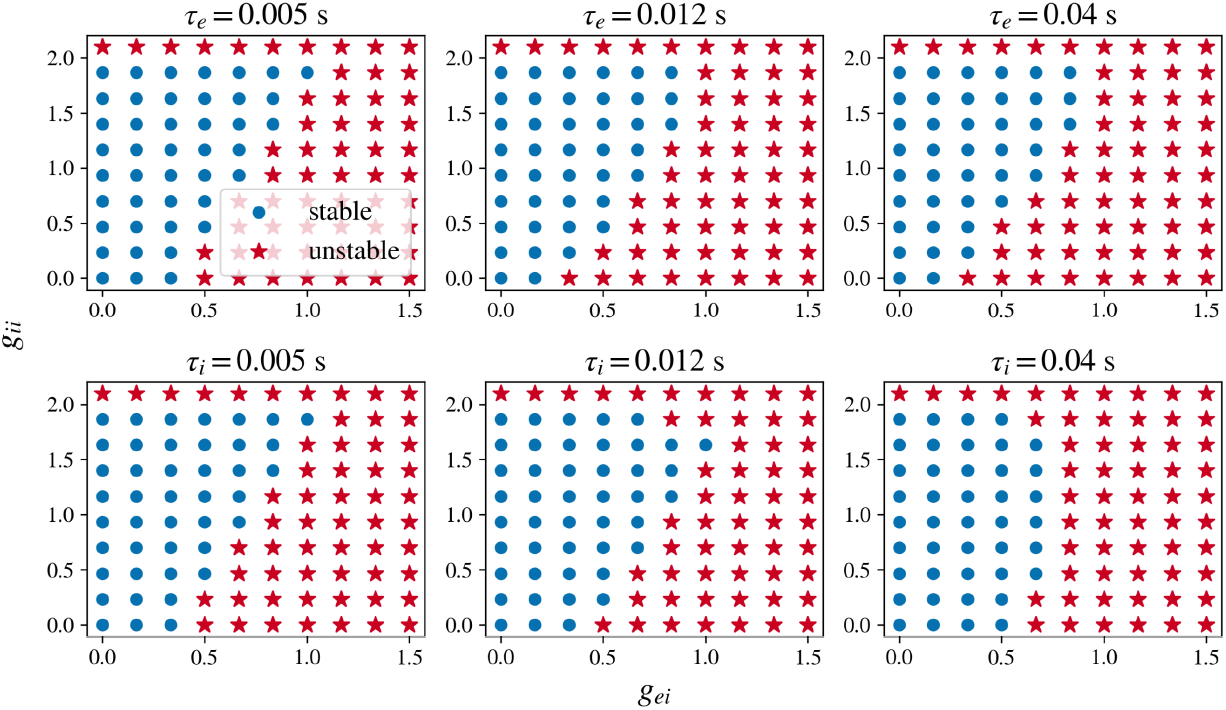
Stability regime for mesoscopic model. Stability regimes plotted for different values of *τ*_e_ and *τ*_i_. When only *τ*_e_ is varied, *τ*_i_ is fixed at 0.003. When only *τ*_i_ is varied, *τ*_e_ is fixed at 0.012. Blue dots represent stable parameter combinations and red stars represent unstable ones.

**Figure S4:**
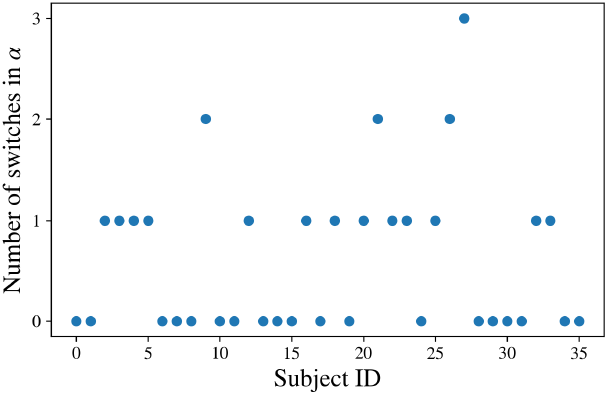
Number of switches in *α*, corresponding to the changes in segregation and integration. Here, the upper bound on *α* is 1. Switches were observed for 17 out of 36 subjects.

**Figure S5:**
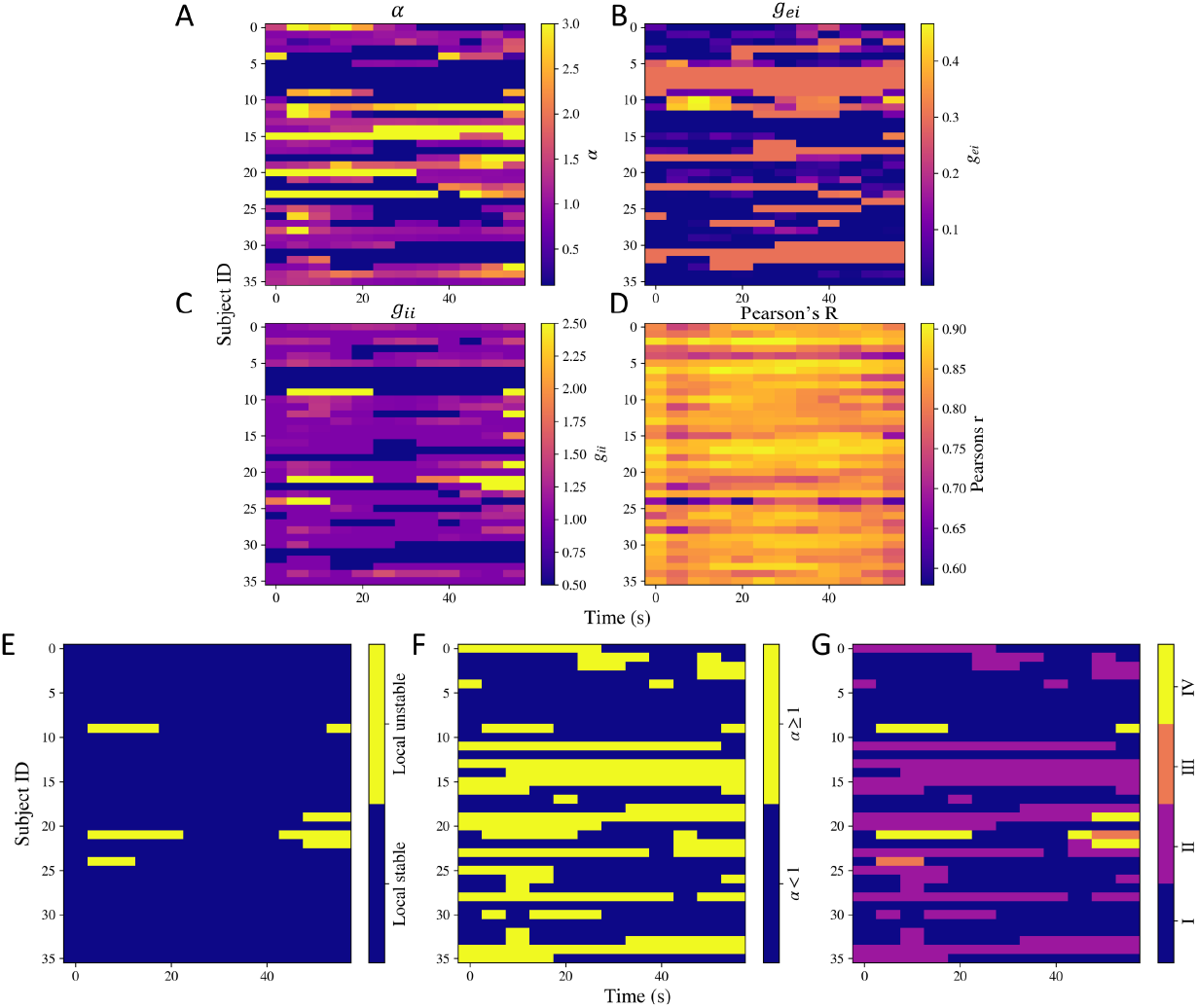
Estimated model parameters **A:** *α*, **B:** *g*_ei_, **C:** *g*_ii_, and **D:** the goodness of fit Pearson’s R calculated at different time points for all the subjects, keeping *α* = 3 as the upper bound. **E, F, G: Dynamic stability. E:** Stability of the local model over time. **F:** Switches in *α* over time. **G:** Switches in different regimes of stability over time. The shade is based on 4 situations: I) Both local model is stable and *α* < 1, II) local model is stable but *α* ≥ 1, III) local model is unstable but *α* < 1, IV) local model is unstable and *α* ≥ 1.

**Figure S6:**
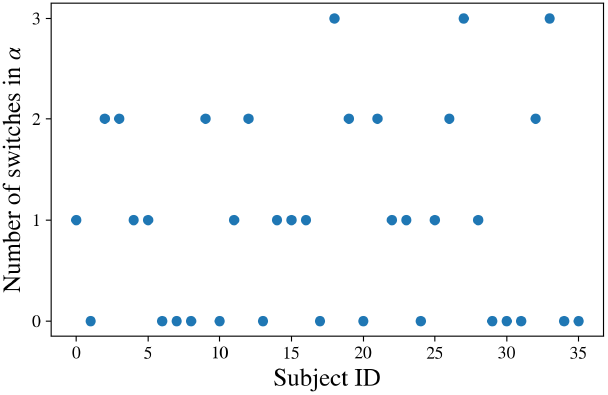
Number of switches in *α*, keeping the upper bound of *α* at 3. The switches correspond to changes in segregation and integration. Switches were observed for 22 out of 36 subjects.

**Figure S7:**
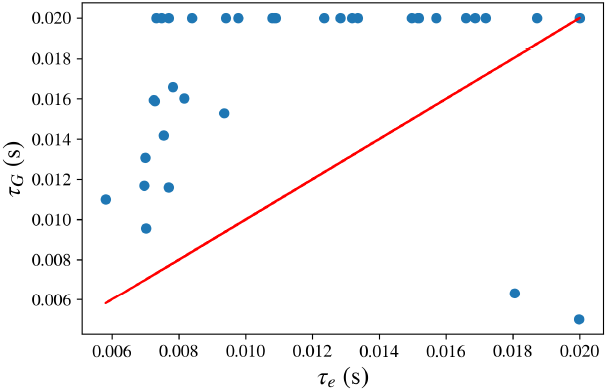
Estimated time constants for the static spectra *τ*_G_ versus *τ*_e_. The red line corresponds to the diagonal line and every point corresponds to a subject. For two subjects, *τ*_G_ was much lower, implying the system is unstable.

## Notes

### Competing Interest Statement

The authors have declared no competing interest.

